# Eastern oyster larvae assemble a core bacterial microbiome distinct from hatchery water treatment systems

**DOI:** 10.64898/2026.01.13.699297

**Authors:** Steph Smith, Ami Wilbur, Rachel T. Noble

## Abstract

Eastern oyster (*Crassostrea virginica*) larvae undergo rapid microbial colonization during early development, a process that may influence larval growth, performance, and hatchery production outcomes. However, the sources and assembly mechanisms of larval-associated bacterial communities remain poorly understood, limiting evidence-based strategies for managing beneficial microbiomes in hatchery settings. We characterized bacterial community composition in 5-day post-fertilization *C. virginica* larvae and across hatchery water treatment stages using *16S rRNA* gene V3-V4 amplicon sequencing. Larval microbiomes were statistically distinct from all sampled water sources (PERMANOVA *q ≤ 0.030*), exhibited 1.9-fold lower Shannon diversity, and showed no relationship to sequential water treatment steps, which themselves did not differ in bacterial composition (all pairwise *q = 0.107*). We identified 39 core bacterial genera present in ≥70% of larval samples, collectively representing 90% of larval sequences and dominated by Rhodobacteraceae/Paracoccaceae (39.5% abundance) and Alteromonadaceae/Marinomonadaceae (24.1%). Critically, 46.2% of core larval sequences belonged to genera highly abundant in larvae but rare (< 0.01%) in all water sources, indicating selective recruitment rather than passive environmental acquisition. Core larval microbiome composition closely parallels that of other marine invertebrates, suggesting conserved host-associated enrichment patterns during early life stages. These results indicate that host-mediated selection dominates larval oyster microbiome assembly, with implications for hatchery management strategies focused on promoting beneficial microbial functions and targeted community supplementation to support consistent larval growth and production efficiency.

**IMPORTANCE:** Eastern oysters generate over $200 million annually in U.S. aquaculture and provide critical ecosystem services including water filtration and coastal habitat formation. Successful hatchery production depends not only on preventing disease and mortality, but also on promoting consistent larval growth, development, and yield. Microbial colonization during early larval stages is increasingly recognized as an important contributor to these outcomes, yet the origins and assembly of beneficial larval microbiomes remain poorly understood. Here, we show that oyster larvae do not simply reflect the microbial composition of hatchery water, but instead actively select a consistent set of bacterial partners in early development. This core larval microbiome is dominated by bacterial families associated with antimicrobial activity, nutrient provisioning, and host growth in other marine systems. Notably, these same bacterial families occur in the larvae of distantly related marine invertebrates, suggesting conserved and potentially beneficial host-microbe associations. Our findings indicate that hatchery water treatment alone is unlikely to determine larval microbiome composition. Instead, targeted strategies that promote or supplement beneficial larval-associated bacteria during early development may provide a more effective path toward improving growth, performance, stability, and overall production efficiency in oyster hatcheries.

## INTRODUCTION

The symbiotic microbial communities associated with marine invertebrates play critical roles in host nutrition, immunity, and development (Yoon et al. 2026). These microbiomes are shaped by complex interactions between environmental conditions, host physiology, and microbial ecology, with implications for organismal fitness and resilience (Yoon et al. 2026). In bivalve mollusks, microbial partners influence digestive efficiency, pathogen resistance, and larval settlement success, making microbiome composition particularly relevant for aquaculture production. Understanding how these microbial assemblages form and function represents a fundamental challenge in marine ecology with direct applications to sustainable shellfish aquaculture.

Adult bivalve microbiomes exhibit host-specific patterns, with core bacterial lineages consistently detected across individuals and populations (Tinning et al. 2025). These core taxa are hypothesized to provide essential functions for host health and development, though their specific roles often remain unclear (Tinning et al. 2025). In adult oysters, taxonomic surveys have revealed microbiome composition varies with tissue type, environmental conditions, and geographic location (Pimentel et al. 2021). In larval oysters, families such as Rhodobacteraceae and Alteromonadaceae have been detected among core microbiome members, often alongside Flavobacteriaceae (Arfken et al. 2021), suggesting these groups are common early symbionts during larval development. Members of these lineages have also been identified in the early life microbiomes of other marine invertebrates, including sea cucumbers (*Apostichopus japonicus*), where they comprise part of a conserved early-life core (Yu et al. 2022). This phylogenetic pattern across distantly related invertebrates suggests that specific bacterial lineages are repeatedly enriched during early larval development, consistent with conserved ecological filtering or functional compatibility with marine invertebrate hosts, rather than obligate host-microbe specificity.

Microbiome assembly during early life stages represents a critical developmental window, as initial colonization events can have lasting effects on host health and performance. In marine invertebrate larvae, microbiome establishment occurs primarily, but not exclusively, through horizontal acquisition from the surrounding environment rather than vertical transmission from parents (Tignat-Perrier et al. 2025).

However, larval microbiomes are not simply passive reflections of environmental bacterial communities. Instead, host immune systems actively shape microbial colonization through pattern recognition receptors (PRRs) that discriminate between beneficial and potentially harmful bacteria (Wang et al. 2019; Yang et al. 2025). In the Pacific oyster *Crassostrea gigas*, fibrinogen-related proteins function as PRRs that recognize specific bacterial surface molecules and mediate selective phagocytosis of particular bacterial taxa, providing a mechanistic basis for host-mediated microbial selection (Wang et al. 2019).

Evidence for selective colonization during early development extends beyond oysters. In white shrimp (*Litopenaeus vannamei*) larvae, Rhodobacteraceae are selectively recruited from rearing water through deterministic assembly processes rather than neutral colonization (Wang et al. 2020). Consistent with these patterns, eastern oyster larvae mount distinct transcriptional responses to probiotic versus pathogenic bacteria, indicating functional immune discrimination early in development (Modak and Gomez-Chiarri 2020). Together, these findings support a model of host-mediated selective colonization during larval development, in which specific bacterial lineages are preferentially recruited rather than passively acquired from the environment.

Beyond community composition, microbial functional plasticity represents a critical frontier for hatchery management, as stable taxonomic assemblages may exhibit variable metabolic activity under changing environmental conditions. For instance, temperature fluctuations may not eliminate core bacterial taxa but can alter production of bioactive compounds, competitive interactions, or nutritional contributions to the host. This functional plasticity suggests that optimizing rearing conditions to promote beneficial microbial activities may be key for improving larval performance and growth. Understanding both the establishment of core larval microbiomes and the environmental factors that modulate their activities will enable more sophisticated management strategies that link microbial ecology to measurable hatchery performance outcomes.

The eastern oyster *C. virginica* represents a species of major aquaculture importance along the Atlantic and Gulf coasts of North America, with production valued at over $200 million annually and significant ecological roles in coastal ecosystems. Understanding larval microbiome dynamics in this species has both fundamental biological significance and direct applications for hatchery management and disease mitigation.

Microbial management in bivalve hatcheries has become increasingly important due to recurring disease outbreaks that threaten production (Petton et al. 2021). Pacific Oyster Mortality Syndrome (POMS) exemplifies these challenges, where viral infection triggers immune dysregulation, microbiome dysbiosis, and opportunistic pathogen colonization (Petton et al. 2021). Interventions targeting specific bacterial taxa have shown promise; bacteriophage therapy reduced larval mortalities by over 90% in both *C. virginica* and *C. gigas* hatcheries through selective elimination of pathogenic *Vibrio* species (Richards et al. 2021). Beyond preventing disease, consistent establishment of beneficial microbiomes may contribute to predictable larval growth trajectories and production efficiency in hatchery systems.

Water treatment systems employed in commercial hatcheries – including filtration, UV sterilization, and setting – substantially alter bacterial community composition in rearing environments (Paralika and Makridis 2025), yet how these treatments influence larval microbiome community assembly remains poorly characterized. Previous studies demonstrated that hatchery-specific environmental conditions influence eastern oyster larval microbiome composition across different facilities (Arfken et al. 2021). However, whether larvae assemble distinct microbiomes from the immediate microbial pool available within a single hatchery system, and whether sequential water treatment steps meaningfully alter that pool, has not been explicitly tested.

Despite growing recognition of microbiome importance in bivalve health and development, fundamental questions remain unresolved. Key questions remain: how larvae selectively acquire beneficial bacteria is not fully understood (Petton et al. 2021), and whether hatchery water treatment practices, which are designed primarily for pathogen reduction, inadvertently impact beneficial bacterial colonization of larvae has received limited investigation. Most microbiome research in oysters has focused on adult life stages (Pimentel et al. 2021) or has examined larvae under controlled laboratory systems rather than in operational hatchery systems where environmental and microbial factors interact with production protocols.

Here, we characterize bacterial communities associated with eastern oyster (*C. virginica*) larvae within a commercial-scale hatchery system and compare them to potential microbial sources in hatchery water. Using *16S rRNA* gene amplicon sequencing, we aim to (1) identify core bacterial taxa consistently associated with oyster larvae, (2) determine whether larval microbiomes differ compositionally from surrounding water sources, and (3) assess how sequential water treatment processes influence bacterial community structure in the hatchery environment. Our findings provide insight into selective colonization processes that shape larval microbiomes and have implications for optimizing microbial management practices.

## RESULTS

### Sample collection and sequencing summary

We collected 22 samples from the UNCW Shellfish Research Hatchery, including seven pooled replicates of larval samples at 5 days post-fertilization, and triplicate water samples from five locations along the water treatment pathway: ambient seawater at the SRH dock, intake tank (60µm filtration), setting tank (5µm filtration), larval tank (0.1µm filtration, UV treatment), and algal culture (used to feed larvae) (Figure 1). *16S rRNA* gene V3-V4 amplicon sequencing yielded 220,687 – 640,145 quality-filtered reads per sample (mean 461,268 ± 134,472 SD; Table S1). After rarefaction to 75,000 reads per sample to normalize sequencing depth, we detected 5,378 amplicon sequence variants (ASVs) across all samples.

**Figure 1.**
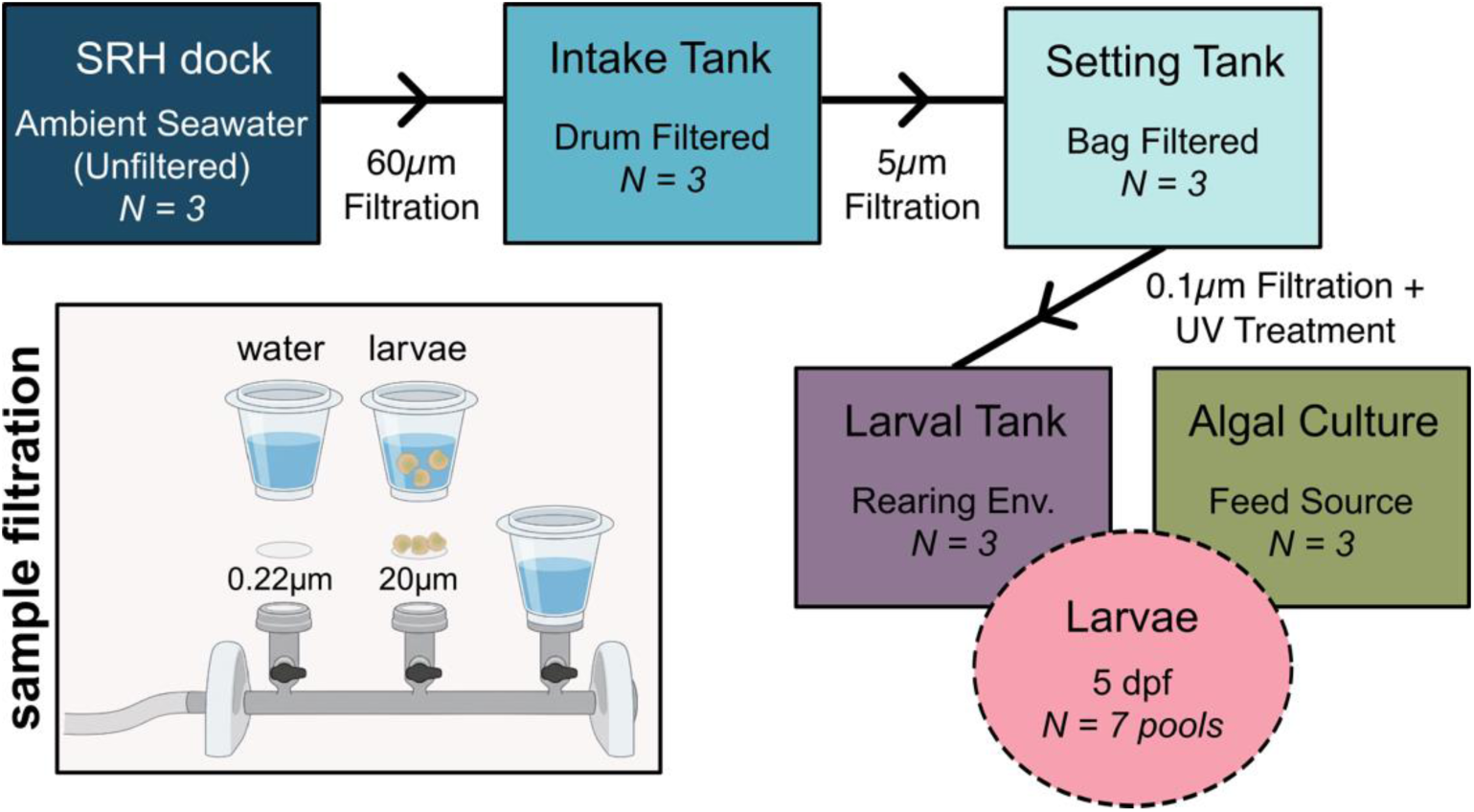
Study system and experimental design. Water treatment pathway showing the progression of seawater through the hatchery system. Ambient seawater from the Shellfish Research Hatchery (SRH) dock undergoes sequential treatment through an intake tank (60µm drum filtration), setting tank (5µm bag filtration), and larval tank (0.1µm cartridge filtration, UV treatment). Algal culture water (used for larval feeding beginning 24h post-fertilization) is maintained separately but exchanges with the larval tank system. Larvae were sampled at 5 days post-fertilization. Inset depicts sample filtration. Water samples were vacuum filtered onto 0.22µm polycarbonate filters; larvae samples were vacuum filtered onto 20µm polycarbonate filters.

### Larval microbiomes are compositionally distinct from all water sources

Principal coordinate analysis (PCoA) of Bray-Curtis dissimilarity revealed that larval microbiomes formed a distinct cluster separated from all water sources along the first principal coordinate axis MDS1, which explained 46.4% of the total variation (Figure 2). Permutational multivariate analysis of variance (PERMANOVA) confirmed that larvae harbored significantly different bacterial communities compared to all five water sources (pseudo-F = 75.80 – 173.00, *q ≤ 0.030* for all pairwise comparisons; Table 1). In contrast, pairwise comparisons among the five water sources revealed no significant differences (*q = 0.107* for all water-water comparisons), despite pseudo-F values ranging from 7.52 to 869.26 (Table 1). The relatively low sample size (n=3 per water source) may have limited statistical power to detect subtle water treatment effects, such that elevated pseudo-F values did not translate into statistically significant differences after multiple-testing correction. Water samples showed substantial variability among MDS2 (24.8% of variation), but this variation did not distinguish among water source types. These results demonstrate that larvae harbor distinct microbial communities compared to their surrounding environment. Sequential water treatment did not produce detectable differences in bulk water bacterial communities, yet larvae remained distinct from all sampled water sources.

**Figure 2.**
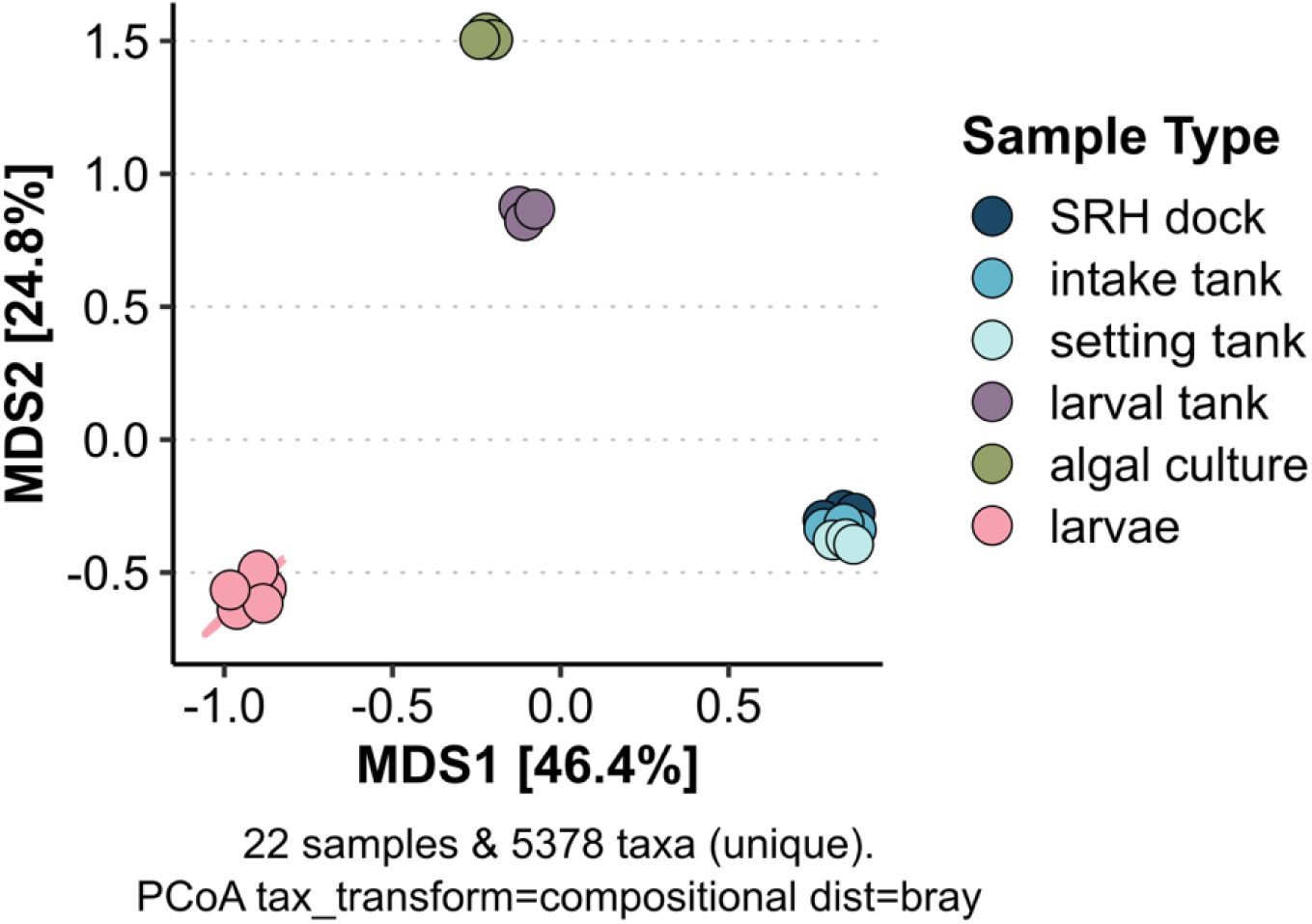
Larvae microbiomes are compositionally distinct from all water sources. Principal coordinate analysis (PCoA) ordination of bacterial community structure based on Bray-Curtis dissimilarity matrices calculated from centered log-ratio (CLR) transformed amplicon sequence variant (ASV) abundances. Each point represents a single sample. Ellipses represents 95% confidence interval around larval group centroid. Analysis based on 5,378 ASVs detected across 22 samples after quality filtering and rarefaction to 75,000 reads per sample.

**Table 1.**
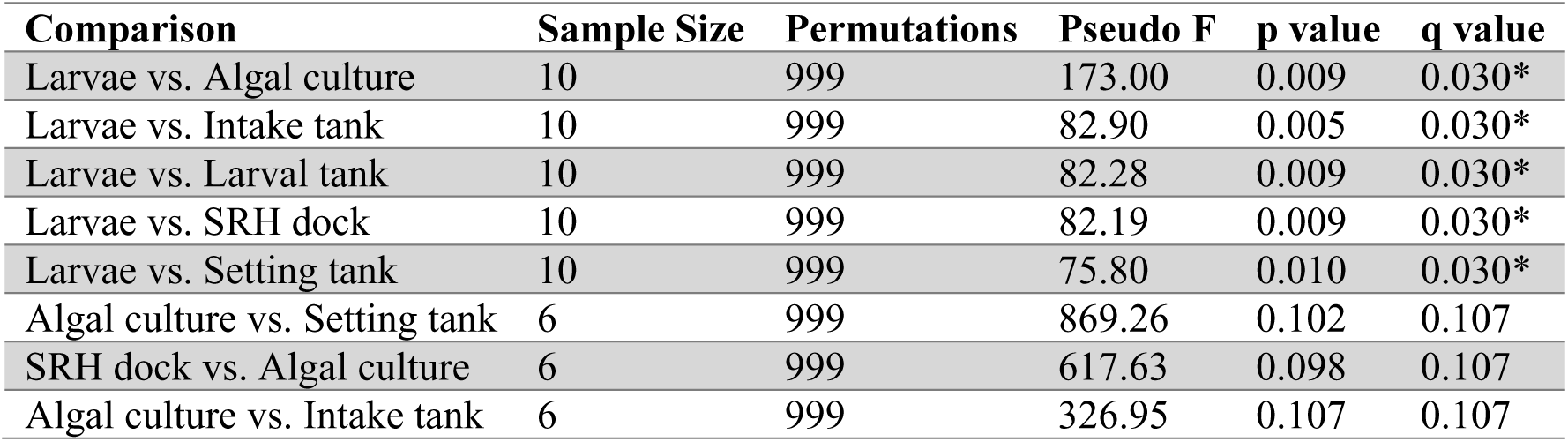
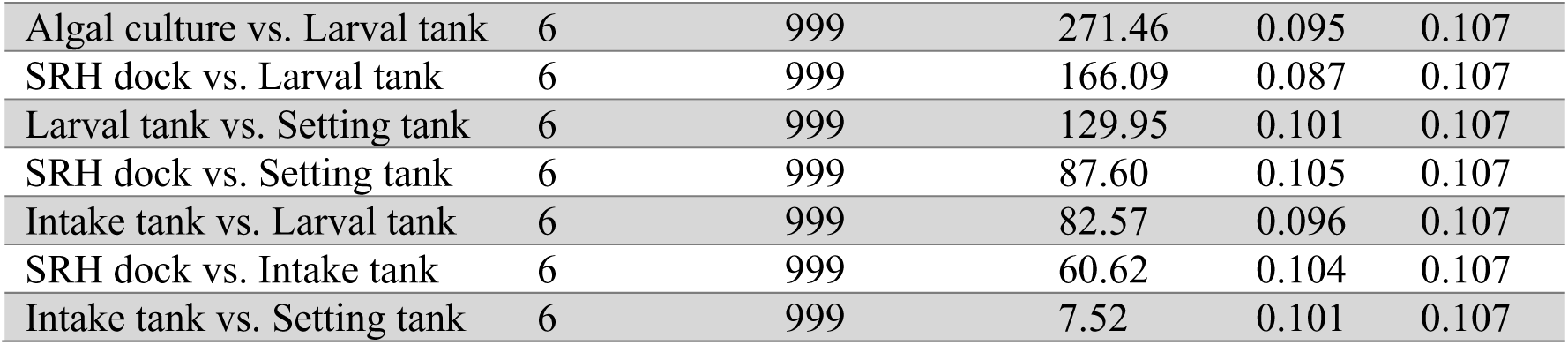
Pairwise PERMANOVA comparisons.

### Oyster larvae exhibit reduced microbial diversity compared to water sources

Larvae exhibited significantly lower Shannon diversity (median = 2.96) compared to all water sources: SRH dock (median = 4.94, *q = 0.017*), intake tank (median = 4.30, *q = 0.017*), setting tank (median = 4.35, *q = 0.017*), and algal culture (median = 2.17, *q = 0.017*) (Figure 3B; Table S2). This represents an approximately 1.9-fold reduction in Shannon diversity in larvae relative to most water sources. Observed ASV richness showed mixed patterns: larvae (median = 1,015 ASVs) had significantly lower richness than SRH dock water (median 3,202, *q = 0.042*) but did not differ significantly from intake tank (*q = 0.517*) or setting tank (*q = 0.333*) (Figure 3A; Table S2). Notably, larval tank water showed the lowest richness among water sources (median = 608 ASVs), which was significantly lower than larvae (*q = 0.056*), while algal culture had the lowest richness overall (median = 91 ASVs, *q = 0.042*). The consistent reduction in Shannon diversity across all water source comparisons indicates that larvae are colonized by a taxonomically restricted subset of bacteria rather than passively accumulating the full diversity present in their rearing environment.

**Figure 3.**
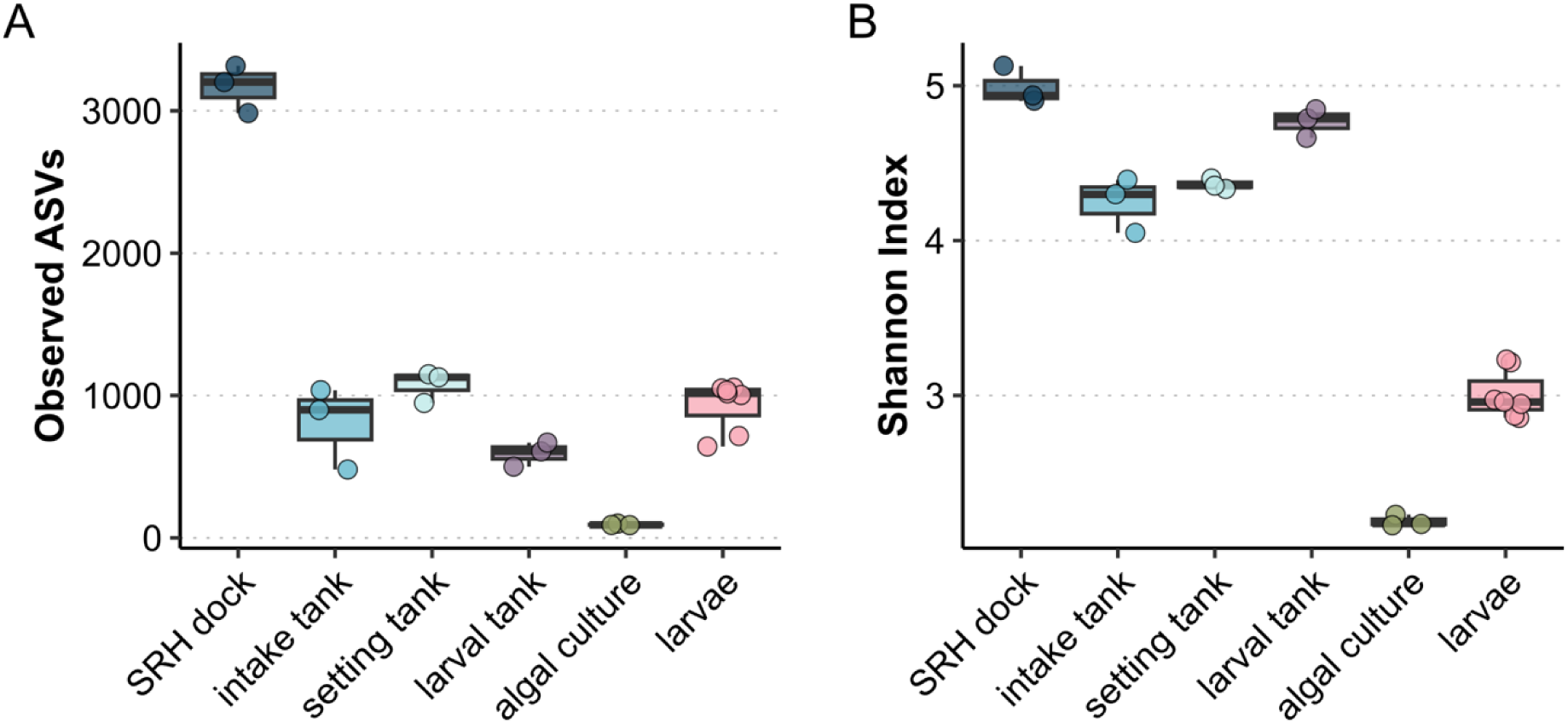
Oyster larvae exhibit reduced microbial diversity compared to surrounding water sources. Alpha diversity metrics comparing bacterial community richness and evenness between larvae and water sources. **(A)** Observed amplicon sequence variants (ASVs). **(B)** Shannon diversity index, which incorporates both richness and evenness. Boxplots show median (center line), interquartile range (box), and full range excluding outliers (whiskers). Statistical comparisons performed on rarefied data (75,000 reads per sample). *q-*values calculated using Benjamini-Hochberg false discovery rate correction. *\*p < 0.05, **q < 0.05*.

### Taxonomic composition reveals family- and genus-level enrichment in larvae

Family-level taxonomic composition analysis revealed distinct patterns between larvae and water sources (Figure 4A). Larval microbiomes were dominated by Rhodobacteraceae (Alphaproteobacteria) and Paracoccaceae (Alphaproteobacteria), which together comprised 39.5% of sequences, and Alteromonadaceae (Gammaproteobacteria) and Marinomonadaceae (Gammaproteobacteria), which together comprised 24.1% of sequences. Cellvibrionaceae (Gammaproteobacteria) represented an additional 4.9% of larval sequences. In contrast, water sources exhibited more variable family-level composition with no single family consistently dominating across all water types.

**Figure 4.**
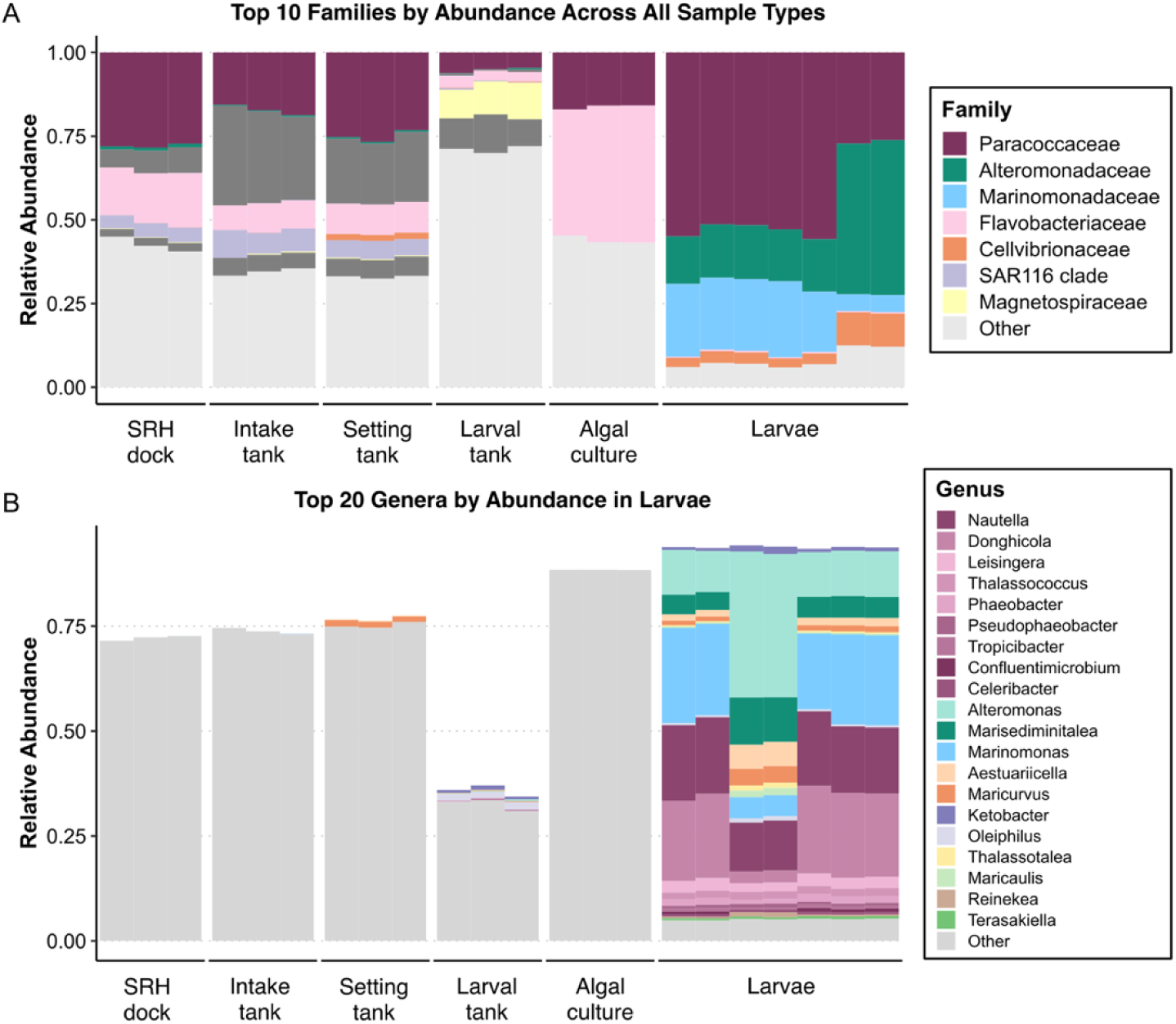
Taxonomic composition reveals family- and genera-level enrichment patterns in larvae. Relative abundance of dominant bacterial taxa across sample types. **(A)** Top 10 most abundant families across all samples. Stacked bar plots show the relative abundance of each family within individual samples, with each bar representing one sample. Taxa not in the top 10 families are grouped as “Other”. **(B)** Top 20 genera detected in larval samples. Bars are colored by family assignment to highlight taxonomic groupings. Taxa not in top 20 genera are grouped as “Other”. All data based on CLR-transformed ASV abundances.

At the genus level, the five most abundant genera in larvae were Alteromonas (Alteromonadaceae, mean 17.4%), Marinomonas (Marinomonadaceae, 16.5%), Nautella (Paracoccaceae, 15.6%), Donghicola (Paracoccaceae, 15.1%), and Marisediminitalea (Alteromonadaceae, 6.5%) (Figure 4B). Together, these five genera comprised 71.1% of larval sequences. The top 20 genera in larvae included ten additional members of Paracoccaceae (Leisingera, Thalassococcus, Phaeobacter, Pseudophaeobacter, Tropicibacter, Confluentimicrobium, Celeribacter, Maritimibacter, Cognatishimia, Marinibacterium, and Pseudooceanicola) and two additional members of Cellvibrionaceae (Aestuariicella and Maricurvus), demonstrating consistent colonization by specific bacterial lineages within these families.

### Core microbiome comprises 39 genera with high prevalence in larvae

Using a 70% prevalence threshold (detected in ≥5 of 7 larval samples) commonly used to identify consistently associated taxa rather than transient colonizers (Shade and Handelsman 2012), we identified 39 genera as core members of the larval microbiome (Figure 5B; Table 2). All 39 core genera were detected in 100% of larval samples except for four genera detected in 71-86% of larvae: Porticoccus (71%), Hyphomonas (86%), Pseudooceanicola (86%), Aquaticitalea (86%), and Hyphomonadaceae Family (86%). The five most abundant genera in larvae, described above, together accounted for 71.1% of sequences in larvae (Table 2). The remaining 34 core genera each comprised 0.12 – 4.7% of larval sequences, collectively representing 28.9% of the core community.

**Figure 5.**
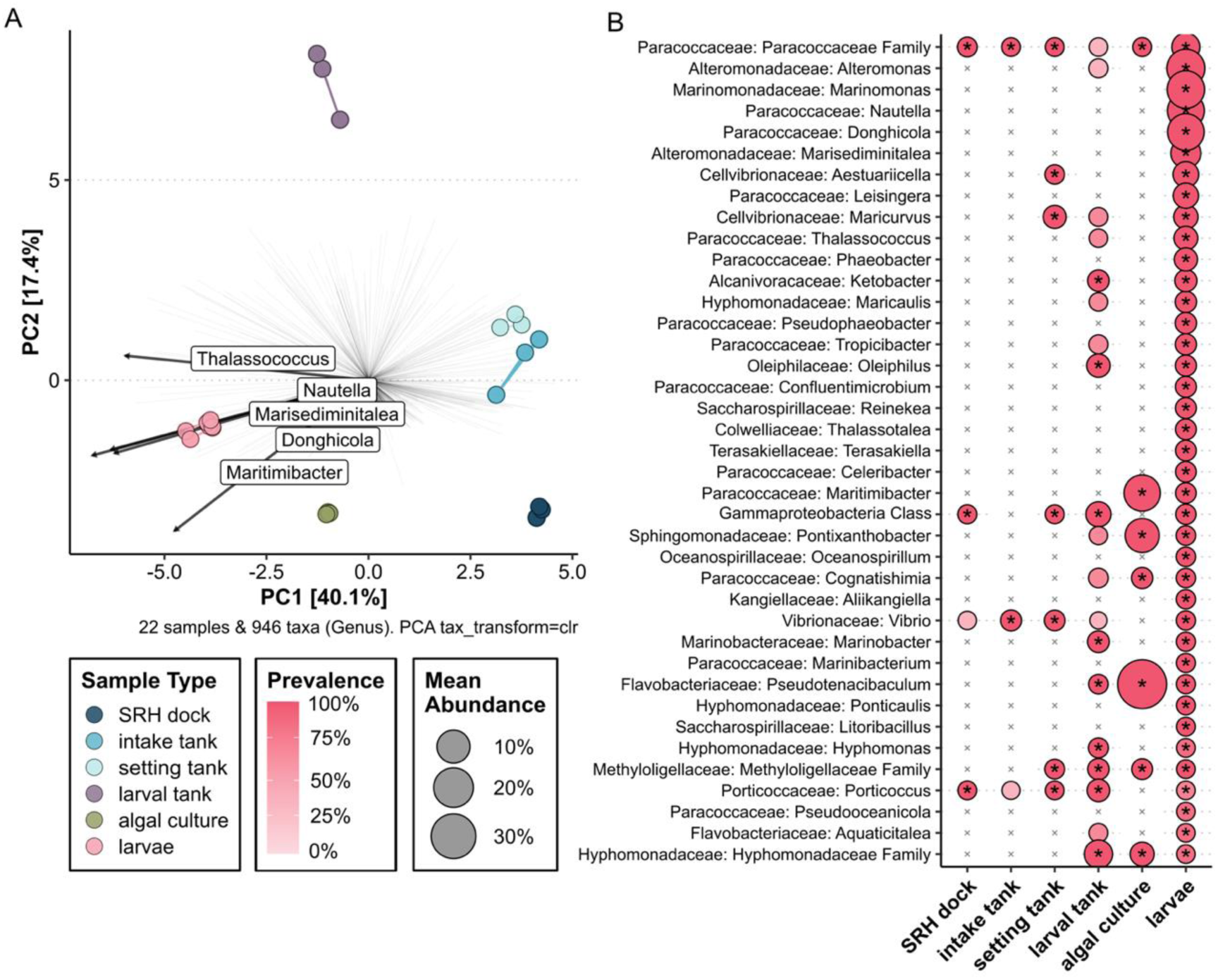
Selective colonization patterns indicate larvae-specific enrichment of core genera. Visualization of selective bacterial colonization in oyster larvae. **(A)** Principal component analysis (PCA) biplot showing genus-level community composition patterns. Arrows represent the top 15 genera contributing to variation along the first two principal components, which explain 48.3% and 18.7% of total variance, respectively. Genera that strongly drive larval separation include *Nautella, Donghicola, Marisediminitalea, Thalassococcus, and Maritimibacter* (arrows pointing toward larvae cluster), while genera associated with water sources point in opposite directions). **(B)** Prevalence heatmap showing detection of the 39 core larval genera (present in 100% of larval samples, n=7/7) across different sample types. Each column represents one sample type with detection frequency indicated by color intensity: dark pink = detected in all replicates (100%), light pink = detected in some replicates, *x* = not detected. Analysis based on presence/absence of genera after quality filtering and taxonomic assignment to the genus level.

**Table 2.**
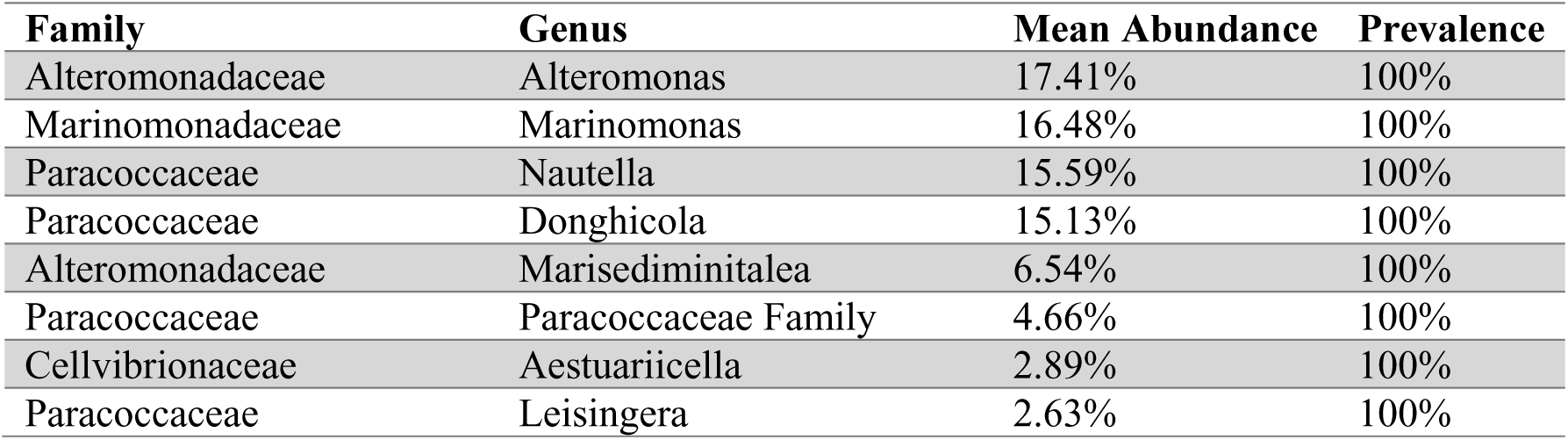
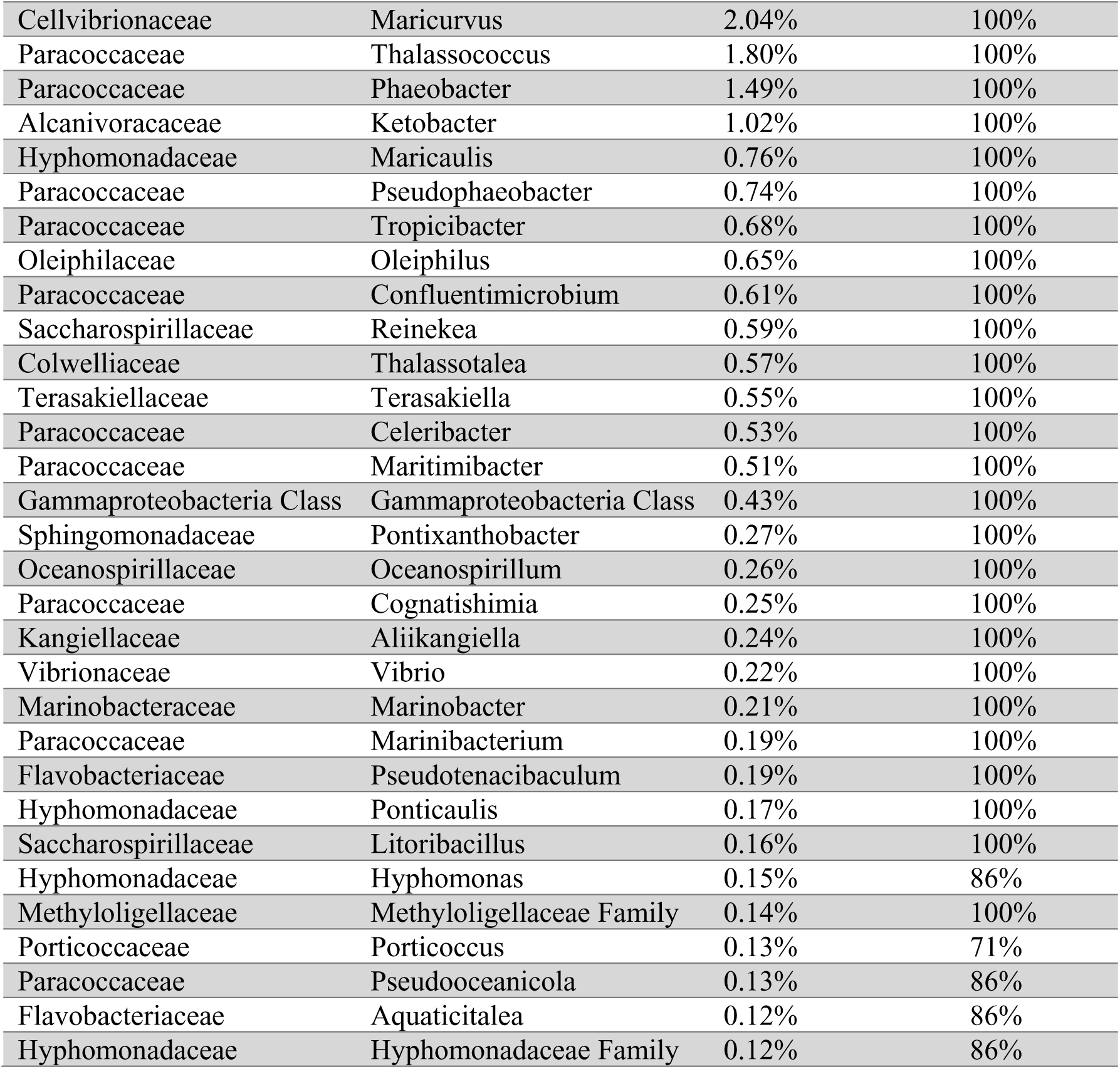
Core bacterial genera in oyster larvae.

Core genera were taxonomically concentrated within Alphaproteobacteria and Gammaproteobacteria. Within Alphaproteobacteria, Paracoccaceae dominated with 14 core genera. Within Gammaproteobacteria, Alteromonadaceae (2 genera), Marinomonadaceae (1 genus), and Cellvibrionaceae (2 genera) were the most abundant families, while additional core genera belonged to Alcanivoracaceae, Oleiphilaceae, Saccharospirillaceae, Colwelliaceae, Kangiellaceae, Vibrionaceae, and Marinobacteraceae (Table 2). This taxonomic composition indicates that larval colonization is dominated by specific lineages rather than representing a random subset of environment bacteria.

### Selective colonization: Core genera are enriched in larvae relative to water sources

Principal component analysis (PCA) of genus-level composition confirmed that larvae-enriched genera drove the separation between larvae and water sources observed in the PCoA analysis (Figure 5A). The first two principal components explained 48.3% and 18.7% of variance, respectively. Genera strongly associated with larvae – including Nautella, Donghicola, Marisediminitalea, Thalassococcus, and Maritimibacter – point toward the larval cluster, while genera associated with water sources point in opposite direction along PC1.

Analysis of core genus detection patterns across sample types revealed selective enrichment in larvae rather than passive accumulation from water sources (Figure 5B; Figure S1; Table S3; Table S4). Of the 39 core larval genera, 18 (46.2%) were classified as “unique to larvae” – detected at high prevalence (≥70%) in larvae but rare or absent (prevalence <30%) in all water sources (Table S3). An additional 14 genera (35.9%) were detected primarily in rearing-associated water sources (larval tank and/or algal culture) but not in external water sources (SRH dock, intake tank, or setting tank). Only 6 genera (15.4%) were found across both rearing-associated and external water sources, and a single genus (2.6%) was detected in external but not rearing water sources (Table S4).

Even when core larval genera were detected in water sources, their prevalence was substantially lower than in larvae (Figure 5B; Figure S1). Of the 39 core genera, only 4 were detected in SRH dock water (mean prevalence 8.5% within that sample type), 3 in intake tank (6.0%), 7 in setting tank (17.9%), 19 in larval tank (37.6%), and 7 in algal culture (17.9%), compared to 100% detection of all 39 genera in larvae (Table S4). Furthermore, only 3 genera were core (≥70% prevalence) within SRH dock samples, 2 in intake tank, 7 in setting tank, 9 in larval tank, and 7 in algal culture, compared to all 39 being core genera in larvae (Table S4). These patterns indicate that larvae selectively enrich specific bacterial lineages rather than passively accumulating bacteria proportional to their availability in the surrounding water.

## DISCUSSION

### Larvae selectively assemble microbiomes that are compositionally distinct from surrounding water sources

Our results demonstrate that five-day post-fertilization eastern oyster larvae assemble microbiomes that are compositionally distinct from all surrounding hatchery water sources, supporting a model of active host-mediated selection during early development.

The observation that larvae exhibited 1.9-fold lower Shannon diversity compared to environmental water sources – despite continuous exposure to high bacterial diversity throughout the water treatment system – provides compelling evidence for active host-mediated colonization rather than passive microbial acquisition. While reduced diversity alone does not prove selection, its consistency across all water source comparisons strongly argues against passive accumulation as the dominant assembly mechanism. This pattern of reduced diversity in early life stages appears consistent across bivalve species, though the direction of change varies developmentally. In Pacific oysters (*Crassostrea gigas*) and related species (*C. corteziensis, C. sikamea*), postlarvae exhibited significantly higher bacterial diversity and richness compared to adults of the same species, with microbiome composition differing substantially between life stages (Trabal Fernández et al. 2014). Developmental stage-specific patterns extend beyond diversity metrics to taxonomic composition. Sydney rock oyster (*Saccostrea glomerata*) larvae exhibit microbiome shifts from D-veliger through pediveliger stages that differ substantially from egg microbiomes despite both being exposed to the same hatchery water, indicating active host-mediated selection operating throughout early development (Unzueta-Martínez et al. 2022). Similarly, eastern oyster larvae across four Chesapeake Bay hatcheries harbored distinct microbiomes that differed by facility despite being reared under comparable protocols, suggesting strong hatchery-specific environmental influences on microbiome assembly (Arfken et al. 2021).

This pattern aligns with progressive niche occupation observed during Pacific oyster development, where diversity variability decreased as core taxa were established (Cho et al. 2024). Pacific white shrimp (*Litopenaeus vannamei*) larvae exhibited selective enrichment of Rhodobacteraceae from rearing water following mouth opening through deterministic assembly processes characterized by positive host selection rather than neutral colonization patterns (Wang et al. 2020). The lower diversity we observed in 5-day post-fertilization larvae relative to water sources may represent an early deterministic selection phase that precedes the higher diversity observed in later-stage postlarvae by Trabal Fernandez et al. (2014), suggesting dynamic microbiome assembly trajectories during early development.

Water treatment did not alter larval microbiome composition. The statistical independence of larval microbiomes from all water sources (PERMANOVA *q ≤ 0.030* for all comparisons), coupled with a lack of differentiation among water sources themselves (all pairwise *q = 0.107*), demonstrates that sequential water treatment steps do not substantially alter bacterial community composition in ways that impact larval colonization. This finding contrasts with documented effects of hatchery water treatment on microbial community structure in marine finfish larviculture, where filtration and UV sterilization can significantly shift bacterial assemblages in rearing water (Paralika and Makridis 2025). The lack of treatment effects in our system may reflect either (1) the specific treatment methods employed at SRH, which does not include chemical treatments, (2) rapid recolonization of treated water from biofilms and air contact, or (3) the dominant influence of host selection masking subtle environmental differences. Regardless, this finding suggests that interventions targeting microbial water quality upstream of larvae may have limited impact on larval-associated bacterial communities, whereas direct modulation of larval microbiomes through probiotic supplementation or selective enrichment of beneficial taxa may prove more effective (Yeh et al. 2020; Richards et al. 2021). While Arfken et al. (2021) demonstrated that hatchery facility itself influences eastern oyster larval microbiome composition when comparing across different hatcheries, our finding that water treatment stages within a single facility show no differentiation suggests that within-facility management of water microbiology has limited influence compared to the strong host-selective processes and facility-specific environmental factors (e.g., biofilms, air exposure, infrastructure) that shape available microbial communities. These conclusions apply specifically to the treatment regine and operational conditions examined here and may differ in systems employing chemical treatments or closed-loop recirculation.

The mechanisms underlying selective colonization likely involve multiple host-mediated processes. Oyster larvae begin feeding on microalgae (algal culture) within 24-48 hours post-fertilization and ingest bacteria alongside or attached to algal cells, potentially explaining the partial overlap between larval and algal culture microbiomes we observed. However, the dramatic enrichment of specific bacterial lineages in larvae (18 genera comprising 46.2% of core sequences that were rare [<0.01%] in all water sources) indicates additional selective mechanisms beyond passive ingestion.

The classification of 18 genera as ‘larvae-unique’ (detected at ≥70% prevalence in larvae but rare [<0.01%] in all water sources) presents two non-mutually exclusive possibilities for their origin. First, these taxa may represent rare environmental bacteria below our detection threshold in water samples that larvae selectively concentrate through active recruitment. Second, they may originate from unsampled sources including broodstock microbiomes through vertical transmission via eggs or sperm. Vertical transmission has been documented in bivalves: pathogenic *Vibrio tubiashii* was detected in carpet shell clam eggs during hatchery mortality events (Dubert et al. 2017), demonstrating that broodstock-associated bacteria can colonize eggs. Similarly, Sydney rock oyster embryos harbored distinct microbiomes from surrounding water, suggesting vertical or maternal provisioning, while progressive shifts through larval development indicated substantial horizontal acquisition (Unzueta-Martínez et al. 2022). The lack of broodstock sampling in our study prevents us from distinguishing between selective recruitment of rare environmental taxa and vertical transmission from parental microbiomes. Both mechanisms are biologically plausible and non-mutually exclusive, and determining their relative contributions will require experimental crosses using broodstock with characterized microbiomes. Understanding these transmission routes has practical implications: broodstock conditioning based on their associated microbial communities could enhance larval microbiome quality through vertical transmission, while manipulation of rearing water or algal cultures could promote beneficial horizontal colonization during early development.

Host immune recognition systems likely play a central role in this selective colonization. Pattern recognition receptors (PRRs) in oysters can discriminate between different bacterial taxa and mediate selective phagocytosis. For example, *C. gigas* fibrinogen-related proteins (FREPs) recognize specific bacterial surface molecules and enhance phagocytic uptake of certain taxa over others (Yang et al. 2021). Similarly, in the bivalve giant clam *Tridacna crocea*, C1q domain-containing proteins exhibit dual functionality, binding both beneficial dinoflagellate symbionts and pathogen-associated molecular patterns, potentially enabling discrimination between beneficial and harmful colonizers (Yi et al. 2025). Critically, *C. virginica* larvae demonstrate functional immune discrimination at molecular and transcriptional levels, mounting distinct responses to probiotic versus pathogenic bacteria, with beneficial bacteria enhancing immune readiness without triggering inflammation while pathogenic bacteria dysregulate immune pathways (Modak and Gomez-Chiarri 2020). These findings indicate that even early-stage bivalve larvae possess immune recognition systems capable of discriminating among bacterial taxa, providing a mechanistic basis for the host-mediated colonization patterns we observed. The rapid establishment of a highly consistent core microbiome by 5 days post-fertilization in our study aligns with this model of active immune-mediated selection, though whether the specific taxa enriched in our larvae (e.g., *Nautella, Donghicola*) are recognized through similar PRR-based mechanisms requires direct investigation.

### Core microbiome dominated by families with documented beneficial functions

The concept of a “core microbiome” – defined as bacterial taxa consistently present across individuals and potentially persistent over time” – has been increasingly applied to oyster species to identify microbial partners likely providing essential host functions (Tinning et al. 2025). In our study, 39 genera detected at ≥70% prevalence in larvae represented such a core community, collectively accounting for 90% of larval sequences. The larval core microbiome was dominated by Rhodobacteraceae/Paracoccaceae (14 genera, 39.5% combined abundance) and Alteromonadaceae/Marinomonadaceae (3 genera, 24.1%). This pattern parallels findings from eastern oyster larvae across multiple Chesapeake Bay hatcheries, where core microbiomes were similarly dominated by Rhodobacteraceae, Flavobacteriaceae, and Alteromonadaceae (Arfken et al. 2021), and from Pacific oyster (*C. gigas*) spat where Rhodobacteraceae consistently dominated across developmental stages and geographic locations (Cho et al. 2024), suggesting these bacterial families provide conserved functions across oyster species and life stages.

This taxonomic pattern is remarkably consistent with findings from sea cucumber (*Apostichopus japonicus*) early life stages, where Alteromonadaceae and Rhodobacteraceae were identified as core microbiome members across six developmental stages spanning three years (Yu et al. 2022). In that system, specific Rhodobacteraceae strains (e.g., *Sulfitobacter* BL28) exhibited growth-promoting effects on larvae, suggesting functional significance for these conserved associations (Yu et al. 2023).

Echinoderms and mollusks are separated by over 500 million years of evolution; the phylogenetic conservation of these bacterial families in early life stages of distantly related marine invertebrates suggests that Alteromonadaceae and Rhodobacteraceae possess traits that are repeatedly favored during larval colonization and are consistent with beneficial functional roles observed in other marine invertebrate systems.

The cross-species conservation of Rhodobacteraceae dominance is particularly striking: Pacific oyster spat harbor Rhodobacteraceae-dominated core microbiomes across developmental stages and geographic locations (Cho et al. 2024), and Rhodobacteraceae are consistently identified as core members in healthy marine invertebrate systems ranging from coral reefs to seagrass beds (Kopprio et al. 2021), suggesting these bacterial families play conserved roles in host development.

The dominance of Rhodobacteraceae in larvae contrasts with adult eastern oyster tissue microbiomes, where the gut is dominated by Mollicutes, with Mollicutes showing significantly lower relative abundance in larvae and biodeposits compared to adult gut tissues (Pimentel et al. 2021). This ontogenetic shift from Rhodobacteraceae-dominated larval communities to Mollicutes-dominated adult gut communities parallels the broader developmental changes observed in Pacific oysters and suggests stage-specific bacterial functions adapted to distinct physiological demands across the oyster life cycle.

Rhodobacteraceae genera identified in our larvae have demonstrated beneficial properties in other marine systems. *Phaeobacter* (1.5% of larval sequences) produces tropodithietic acid with antimicrobial activity against pathogenic *Vibrio* and has been validated as a probiotic in marine fish larvae (Dittmann et al. 2020). *Leisingera* (2.6%) is part of the core bacteriota in healthy shrimp larvae, and *Leisingera* isolates from Hawaiian bobtail squid produce antimicrobial compounds and provide chemical defense in egg jelly coats (Suria et al. 2020). More broadly, the Rhodobacteraceae family maintained overwhelming dominance after mouth opening in *Litopenaeus vannamei* larvae, where they are recruited via positive host selection from rearing water (Wang et al. 2020). Probiotic supplementation that enriched gut Rhodobacteraceae in shrimp demonstrated enhanced digestive enzyme activity and immune responses through upregation of amino acid meatbolism and NF-kB signaling pathways (Ding et al. 2022). While the dominant genera *Nautella* (15.6%) and *Donghicola* (15.1%) lack genus-specific characterization, their family (Rhodobacteraceae) is well-documented for photoheterotrophic metabolism, antimicrobial metabolite production, and beneficial symbioses with marine invertebrates. These multi-faceted functional capabilities, including antimicrobial defense, nutritional provisioning, and immune modulation, may explain the conserved dominance of Rhodobacteraceae across marine invertebrate larvae.

We note that the taxonomic identification from *16S rRNA* gene amplicon sequencing provides compositional but not functional information. While Rhodobacteraceae and related families are known for producing antimicrobial compounds, synthesizing B vitamins, and providing other beneficial functions in various marine systems, whether the specific strains detected in our larvae encode these capabilities requires direct validation. The functional roles we discuss represent hypotheses based on taxonomic affiliation that require experimental confirmation in the eastern oyster larval system.

Alteromonadaceae and the closely related Marinomonadaceae (combined 24.1% of larval sequences) represent the second major component of the core microbiome. Members of these families are recognized as beneficial bacteria in marine invertebrate aquaculture, commonly found in healthy shrimp and bivalves, and have been utilized as probiotics in larviculture systems (Sutanti et al. 2024).

Alteromonadaceae were among the dominant taxa in early D-veliger stage Sydney rock oyster larvae (Unzueta-Martínez et al. 2022), consistent with our findings in 5-day post-fertilization eastern oyster larvae (equivalent to late D-stage), suggesting this family plays conserved early developmental roles across oyster species. *Alteromonas* species often produce bioactive compounds that can inhibit pathogen growth and contribute to larval settlement processes in some marine invertebrates (Weiner et al. 1985), though whether the specific taxa we detected serve similar roles in oyster larvae remains to be demonstrated.

An important limitation is that *16S rRNA* gene amplicon sequencing provides taxonomic identification but cannot directly assess functional capabilities or metabolic activity. The presence of genera within families known for beneficial traits does not confirm that the specific strains in our larvae possess or express these capabilities. The high abundance of poorly characterized genera (*Nautella, Donghicola, Marinomonas*) highlights substantial gaps in our understanding of oyster-associated bacteria and underscores the need for cultivation-based and functional genomic approaches to validate hypothesized beneficial roles.

### Study limitations and future research priorities

Three primary limitations constrain the interpretation of our findings. First, we sampled from a single hatchery under standard operating conditions (28°C, ∼30 ppt salinity). Environmental factors including temperature, salinity, pH, and food availability can substantially influence bivalve microbiomes (Pierce and Ward 2019; Zhong et al. 2024), and our results may not generalize to different rearing conditions or developmental temperatures. Understanding how environmental variation modulates host selection and microbiome assembly – and whether this variation affects larval health outcomes – represents an important avenue for future research with direct implications for optimizing hatchery practices.

Temperature effects on oyster microbiomes are well-documented: heat stress causes microbiome dysbiosis in Pacific oysters with shifts toward opportunistic colonizers (Green et al. 2019; Zhong et al. 2024), while disease-resistant Kumamoto oysters maintain stable, probiotic-enriched microbiomes under pathogen challenge that susceptible oysters lose (Dittmann et al. 2020). These findings suggest that environmental stressors may compromise host-selective processes, allowing opportunistic colonization that our study conducted under ambient hatchery conditions (28°C) may not have captured.

Second, we sampled larvae at a single developmental timepoint (5 days post-fertilization, D-stage). Studies examining multiple developmental stages reveal dynamic microbiome shifts throughout early ontogeny (Unzueta-Martínez et al. 2022; Cho et al. 2024). Although the five-day larval stage represents a transitional developmental period, our findings that larvae harbor fundamentally distinct microbiomes compared to all sampled water sources still raises the critical question of how host-mediated selection operates to establish this compositional divergence. Whether the core microbiome we identified at this stage represents a stable community that persists through settlement or a transitional community that differs from later life stages remains to be determined. Important questions remain regarding assembly dynamics and microbial persistence across life stages, which will be critical for understanding whether larval interventions have lasting effects on juvenile and adult oyster health. Additionally, samples in this study represent pooled larvae from a single spawn event, limiting generalizability across different broodstock genetic backgrounds or spawning conditions.

Third, our source attribution analysis is limited to sampled water sources and cannot definitively identify the origin of larvae-enriched taxa. We did not sample broodstock adults, hatchery tank biofilms, or other yet unidentified reservoirs that may serve as colonization sources. Comprehensive spatial and temporal sampling coupled with experimental manipulation would better resolve colonization sources and mechanisms.

### Implications for hatchery management and aquaculture

Our findings suggest that microbial management strategies in oyster hatcheries should focus on direct larval microbiome modulation rather than solely on water treatment optimization. The statistical independence of larval microbiomes from water treatment stages, combined with evidence for selective host-mediated colonization, indicates that conventional approaches focused on pathogen reduction in water may be insufficient to ensure beneficial microbial colonization of larvae. Instead, strategies that enhance selective enrichment of beneficial taxa, such as probiotic supplementation or broodstock selection based on associated microbiomes, may prove more effective (Yeh et al. 2020). The consistent establishment of Rhodobacteraceae-dominated communities by 5 days post-fertilization suggests this early developmental window may represent a critical period for intervention, though experimental validation is required to assess impacts on larval survival, growth, and disease resistance through later developmental stages.

While our study characterized the taxonomic composition of the larval core microbiome, translating these findings into hatchery management strategies requires understanding how environmental conditions modulate the functional activities of these bacterial communities. The consistent presence of core taxa like Rhodobacteraceae and Alteromonadaceae across diverse oyster systems suggests compositional stability; however, their functional contributions likely vary with temperature, pH, dissolved oxygen, and other abiotic factors. Environmental conditions can modulate microbial functional activities even when community composition remains relatively stable. In marine sponges exposed to naturally acidified waters, the composition of host microbiome was largely conserved between control and CO_2_ seep sites, yet the microbiome demonstrated reduced capacity for carbohydrate update and nitrogen metabolism under acidified conditions (Botté et al. 2019). This functional plasticity suggests that optimizing temperature, pH, dissolved oxygen, and nutrient availability during critical developmental windows could enhance beneficial microbial activities (*e.g.,* antimicrobial compound production by *Phaeobacter* or biofilm formation by Rhodobacteraceae) without necessarily altering community structure.

Beyond traditional approaches to bulk water treatment, several targeted strategies could enable more effective manipulation of larval microbiomes. Bioencapsulation in live feed represents one promising avenue: beneficial bacteria could be encapsulated within or attached to algal cells for co-delivery with nutrition during feeding, leveraging natural feeding behavior to enhance colonization efficiency (Masoomi Dezfooli et al. 2021). Biofilm-based delivery systems offer another approach, where beneficial biofilms on tank surfaces could provide continuous low-level inoculation, stabilizing communities throughout larval development (Gao et al. 2022). Additionally, since vertical transmission may contribute to larval colonization (Unzueta-Martínez et al. 2022), conditioning broodstock with beneficial bacteria could enhance microbial transfer to eggs before exogenous feeding begins. These approaches require hatchery validation but represent tractable engineering solutions that move beyond water treatment toward direct, mechanistic manipulation of larval microbiomes.

This evidence-based approach to environmental manipulation complements compositional characterization and could enable hatchery managers to develop protocols linking specific culture conditions to measurable functional outcomes: reduced mortality through enhanced antimicrobial defenses, optimized growth rates via efficient nutrient cycling, and selection for broodstock whose larval microbiomes maintain robust benficial activities across production cycles and environmental variation.

The efficacy of probiotic supplementation in flow-through hatchery systems depends critically on whether introduced strains match host selection criteria. Our evidence for selective enrichment of specific taxa suggests that larvae may discriminate against non-selected strains regardless of probiotic dosing.

Understanding the molecular basis of host selection represents a critical research priority for rational probiotic design and for translating microbiome ecology into predictive hatchery management strategies.

## METHODS

### Larval oyster culture

Eastern oyster (*C. virginica*) larvae were obtained from the Shellfish Research Hatchery (SRH; University of North Carolina Wilmington, NC, USA) and maintained following standard hatchery procedures.

Larvae were reared in filtered, UV-treated water obtained from the hatchery at 28°C and ∼30 ppt with gentle aeration. Algal diets consisted of a mixed diet (*Chaetoceros mulleri, C. neogracilis, Isochrysis lutea, and Pavlova lutheri*) supplied at rations following Helm et al. 2004.

### Study site and sample collection

Samples were collected on May 9, 2025 from the UNCW Shellfish Research Hatchery (SRH) in Wilmington, North Carolina, USA. Eastern oyster (*Crassostrea virginica*) larvae were spawned approximately 5 days prior to sampling and maintained in a hatchery system supplied with water from the adjacent UNCW dock. Influent seawater undergoes sequential filtration through 60µm, 5µm, and 0.2µm cartridge filters, followed by UV sterilization, before entering larval rearing tanks. Larvae were fed daily with on-site microalgal culture starting 2 days post-fertilization.

Water samples (n=15) were collected from five sources in triplicate: SRH dock water, intake tank (60µm filtration), setting tank (5µm filtration), larval rearing tank (0.2µm filtration, UV sterilization), and algal culture tank. For algal cultures, 150mL whole culture (including algal cells with associated bacteria) was filtered through 0.22µm polycarbonate filters, capturing both algae-associated and planktonic bacteria that larvae would ingest during feeding. For each water sample, 150mL was filtered through 0.22µm polycarbonate filters (Whatman) using vacuum filtration. Larval samples (n=7) represent pooled technical replicates from a single spawn event, with each sample containing larvae collected from 150mL of rearing water maintained at approximately 8 larvae/mL. Larvae were captured by filtering through 20µm polycarbonate filters to selectively capture larvae and physically associated microbes while excluding free-living planktonic bacteria. All filters were stored on ice and transported to the UNC Institute of Marine Sciences (Morehead City, NC) within 4 hours of collection, then stored at -80°C until DNA extraction.

### DNA extraction

Genomic DNA was extracted from all filters using the MagMAX Microbiome Ultra Nucleic Acid Isolation Kit (Thermo Fisher Scientific) on a KingFisher Apex automated extraction system following the manufacturer’s protocol for bacterial DNA isolation. DNA concentrations were quantified using a Qubit 4.0 fluorometer with the dsDNA High Sensitivity Assay Kit (Thermo Fisher Scientific). Extracted DNA was stored at -80°C until sequencing.

### 16S rRNA gene amplicon sequencing

The V3-V4 hypervariable region of the *16S rRNA* gene was amplified using primers 341F (5’-CCTAYGGGNBGCWGCAG-3’) and 805R (5’-GACTACNVGGGTMTCTAATCC-3’). Library preparation and sequencing were performed by SeqCenter (Pittsburgh, PA, USA) on an Illumina NovaSeq platform generating 2x250 bp paired-end reads. Sequencing yielded 220,687 to 640,145 reads per sample (mean = 461,268 ± 134,472 SD). Extraction blanks were sequenced in parallel with experimental samples and yielded negligible read counts, indicating minimal background contamination.

### Bioinformatic processing

Raw sequence data were processed through the nf-core/ampliseq pipeline version 2.15.0 (Straub et al. 2020). Briefly, raw reads were quality filtered and trimmed using Cutadapt v4.6 (Martin 2011) to remove primer sequences and low-quality bases. Amplicon sequence variants (ASVs) were inferred using DADA2 v1.30.0 (Callahan et al. 2016) with default parameters for quality filtering, denoising, and chimera removal. Taxonomy was assigned to ASVs using the SILVA reference database version 138.2 (Quast et al. 2012) with the naïve Bayesian classifier implemented in DADA2. The pipeline generated 7,492 ASVs, which were further filtered to remove chloroplast, mitochondrial, and domain-level unassigned sequences, resulting in 5,378 ASVs for downstream analysis.

### Statistical analyses

All statistical analyses were performed in R version 4.3.2 using the phyloseq package v1.46.0 (McMurdie and Holmes 2013) for data organization and the vegan package (Oksanen et al. 2023) for ecological statistics. Visualizations were generated using ggplot2 (Wickham 2011) and microViz (Barnett et al. 2021).

### Alpha diversity

Alpha diversity was assessed using observed ASV richness and Shannon diversity index, both calculated on rarefied data (75,000 reads per sample) to account for differences in sequencing depth. Rarefaction was performed using the phyloseq rarefy_even_depth function. Statistical differences in alpha diversity across sample types were evaluated using Kruskal-Wallis tests followed by pairwise Wilcoxon rank-sum tests with Benjamini-Hochberg false discovery rate (FDR) correction for multiple comparisons.

### Beta diversity

Beta diversity was assessed using principal coordinate analysis (PCoA) based on Bray-Curtis dissimilarity calculated from centered log-ratio (CLR) transformed ASV abundances. All beta diversity analyses were performed on unrarefied, CLR-transformed data to appropriately handle compositional data. Permutational multivariate analysis of variance (PERMANOVA) with 999 permutations was performed using the adonis2 function in vegan to test for significant differences in community composition among sample types, followed by pairwise PERMANOVA tests with FDR correction. Ordinations were visualized using principal component analysis (PCA) on CLR-transformed genus-level abundances to facilitate taxonomic interpretation.

### Core microbiome and source attribution

Core microbiome composition was determined by identifying taxa present in ≥70% of samples within each sample type. This threshold follows established precedent for defining core microbiomes (Shade and Handelsman 2012). Prevalence categories were further identified as core (≥70%), common (50-69%), present (30-49%), rare (<30%), and undetected. Source attribution analysis was conducted by cross-referencing core and common taxa detected in larvae (prevalence ≥50%) with their detection frequency across all water source types to identify taxa unique to larvae versus those shared with specific water sources.

## Data availability

Raw sequence data and associated metadata have been deposited in the NCBI Sequence Read Archive under BioProject accession PRJNA1400077.

## Authors’ Contributions

Steph Smith: Conceptualization, Methodology, Investigation, Formal Analysis, Writing – Original Draft, Visualization, Project Administration

Ami Wilbur: Resources, Methodology, Writing – Review & Editing

Rachel Noble: Conceptualization, Funding Acquisition, Supervision, Writing – Review & Editing All authors read and approved the final manuscript.

## Acknowledgements

We thank the UNCW Shellfish Research Hatchery staff and students for providing larval oysters and access to the hatchery facilities essential to this study.

